# The intra-individual reliability of ^1^H-MRS measurement in the anterior cingulate cortex across one year

**DOI:** 10.1101/2023.06.07.544020

**Authors:** Meng-Yun Wang, Max Korbmacher, Rune Eikeland, Alexander R. Craven, Karsten Specht

## Abstract

Magnetic resonance spectroscopy (MRS) is the primary method that can measure the levels of metabolites in the brain in vivo. To achieve its potential in clinical usage, the reliability of the measurement requires further articulation. Although there are many studies that investigate the reliability of gamma-aminobutyric acid (GABA), comparatively few studies have investigated the reliability of other brain metabolites, such as glutamate (Glu), N-acetyl-aspartate (NAA), creatine (Cr), phosphocreatine (PCr), or myo-inositol (mI), which all play a significant role in brain development and functions. In addition, previous studies which predominately used only two measurements (two datapoints) failed to provide the details of the time effect (e.g., time-of-day) on MRS measurement within subjects. Therefore, in this study, MRS data located in the anterior cingulate cortex (ACC) were repeatedly recorded across one year leading to at least 25 sessions for each subject with the aim of exploring the variability of other metabolites by using the index coefficient of variability (CV); the smaller the CV, the more reliable the measurements. We found that the metabolites of NAA, tNAA, and tCr showed the smallest CVs (between 1.61 and 4.90 %), and the metabolites of Glu, Glx, mI, and tCho showed modest CVs (between 4.26 and 7.89 %). Furthermore, we found that the concentration reference of the ratio to water with tissue correction results in smaller CVs compared to the ratio to tCr. In addition, we did not find any time-of-day effect on the MRS measurements. Collectively, the results of this study indicate that the MRS measurement is reasonably reliable in quantifying the levels of metabolites.

**Key points:** The MRS measurement is reliable within subject.

The ratio to water provides more reliable results compared to the ratio to tCr.

The ratio to water is recommended as the internal concentration reference.

## 1. Introduction

Magnetic resonance spectroscopy (MRS) can non-invasively quantify the levels of metabolites in the central neural system by measuring different resonance frequencies of the proton hydrogen (^1^H) embedded in them (Oz et al., 2014; Wilson et al., 2019). In typical applications, a 5-minute acquisition time can provide a good-quality dataset, where the spectra of a single voxel can be obtained (Wilson et al., 2019). With MRS, one can distinguish different brain lesions with similar MRI appearance (Oz et al., 2014). In addition, it can be used to investigate the neurometabolic responses to external stimuli in-vivo among healthy and patient populations (Pasanta et al., 2023). Therefore, it is an important means and can be used to assist clinical diagnosis, monitor treatment effects, and facilitate patient management (Oz et al., 2014). However, to fully unleash its potential in investigating neurometabolic responses and to advance its clinical usage, the reliability of the MRS measurement should be articulated.

Indeed, there are several studies focusing on the reliability of MRS with the main focus on the reliability of gamma-aminobutyric acid (GABA), where test-retest analysis was used between sessions (Baeshen et al., 2020; Brix et al., 2017; Duda et al., 2021; Mikkelsen, Singh, Sumner, & Evans, 2016; Near, Ho, Sandberg, Kumaragamage, & Blicher, 2014) or within sessions (Brix et al., 2017; O’Gorman, Michels, Edden, Murdoch, & Martin, 2011). GABA is the principal inhibitory neurotransmitter in the human brain, which is crucial for normal neurological function and plays a significant role in learning, memory, and other cognitive functions (Pasanta et al., 2023). It is shown that the reliability of GABA in the brain, which is indicated by the coefficient of variability (CV), varies between 4%-15% depending on region, quantification techniques and the study design (Baeshen et al., 2020; Brix et al., 2017; Duda et al., 2021; Mikkelsen et al., 2016; Near et al., 2014; O’Gorman et al., 2011). Even in the same brain region, however, the CV can be different. For example, one study showed that the CV of GABA among the anterior cingulate cortex (ACC) was 8% using the ratio to total creatine (tCr) while 7.5% with the ratio to water (Duda et al., 2021).

To be notice, all the aforementioned studies used a 3T MRI scanner and MEGA-PRESS to detect GABA concentration level. Accordingly, the wide range of the CV is not only because of different brain locations but also due to different quantification indices used in these studies. There are two commonly used quantification indices: the ratio to tCr, and the ratio to water with tissue correction (Gasparovic et al., 2006; Near et al., 2021), which will be referred to the ratio to water, for the sake of simplicity. It is suggested that the ratio to water performs better regarding the reliability of the GABA (Duda et al., 2021).

Although much insight has been made regarding GABA, surprisingly, very few studies investigated the reliability of other critical brain metabolites (Kirov et al., 2012; Mullins et al., 2003; Schirmer & Auer, 2000; van Veenendaal et al., 2018), which also play significant roles in brain function. For example, glutamate (Glu) is the most abundant excitatory neurotransmitter, which plays a significant role in brain cognition and neurological development by countering balance with GABA; N-acetyl-aspartate (NAA) and N-acetyl-aspartyl-glutamate (NAAG) are neuromodulators which inhibit the synaptic release of GABA, glutamate, and dopamine, and regulate GABA receptor expression; myo-inositol (mI) is a key precursor of membrane phospho-inositides and phospholipids, and is considered as a glial marker by involving in the cell membrane and myelin sheet structures (Haris, Cai, Singh, Hariharan, & Reddy, 2011; Harris, Saleh, & Edden, 2017).

Besides the limited numbers, the aforementioned studies also possess their own limitations. For example, two studies focused on the white matter (WM) instead of the grey matter (GM) (Mullins et al., 2003; Schirmer & Auer, 2000), of which the participants of one study are only schizophrenia patients (Mullins et al., 2003). One study only focused on Glu (van Veenendaal et al., 2018) while the remaining one covered almost a third of the brain (Kirov et al., 2012) which is a deviation from the most widely used method in MRS named single-voxel spectroscopy (Wilson et al., 2019). Furthermore, none of them investigates the different performance between the ratio to tCr and water on the reliability of the MRS measurement. In addition, previous studies generally have very low sampling rates within subjects, where only two measurements were predominately collected within a time period ranging from the same day to several months (Baeshen et al., 2020; Brix et al., 2017; Duda et al., 2021; Mikkelsen et al., 2016; Mullins et al., 2003; Near et al., 2014; O’Gorman et al., 2011; Schirmer & Auer, 2000; van Veenendaal et al., 2018). The low sampling rate fails to illustrate the details of the time effect on MRS measurements, such as the time-of-day effect manifesting on the functional brain organization (Orban, Kong, Li, Chee, & Yeo, 2020; Vaisvilaite, Hushagen, Gronli, & Specht, 2022).

Therefore, in this study, three subjects were repeatedly scanned in an MRI scanner across one year, resulting in at least 25 MRS sessions for each subject. To be noticed, this study is part of a project named Bergen breakfast scanning club (BBSC) where we also collected structural and functional MRI data (Korbmacher et al., 2023; Wang, Korbmacher, Eikeland, & Specht, 2022, 2023). The aims of the study are threefold. First, to investigate the reliability of other brain metabolites other than GABA. Second, to explore whether the ratio to water can generate more reliable results compared to the ratio to tCr. Third, to explore the time-of-day effect on the levels of the metabolites.

GABA is a weak signal that often requires dedicated spectral editing techniques to be resolved and often would be compromised by artifacts and scanner drift. Given that, we posit that the reliability of other metabolites is better than that of GABA.

## 2. Methods

### 2.1 Participants

Three participants (**Table 1**) were scanned in this study, which is part of a precision brain mapping project titled BBSC Project (Korbmacher et al., 2023; Wang et al., 2022, 2023). In this project, we aim to chart the individual brain organizations and explore the reliability of the MRI data by repeatedly scanning three subjects for a year including resting state fMRI data, MRS data, and structural brain imaging data (Wang et al., 2022, 2023). The participants were repeatedly scanned twice a week between February 2021 and February 2022 except for the summer (Jun. to Sep. 2021) and winter (Dec. 2021 to Jan. 2022) breaks. In total, there are 38, 39, and 25 MRS sessions for subjects 1, 2, and 3, respectively.

**Table 1.** Basic demographic information of subjects

|  | 1 | 2 | 3 |
| --- | --- | --- | --- |
| Gender | Male | Male | Male |
| Age | 31 | 27 | 40 |
| Laterality | Right | Right | Right |
| Regular caffeine consumption? | No | Yes | Yes |
| Regular nicotine consumption? | No | No | No |

They are all male and can speak at least two languages (their native languages, and English). To be noticed, subject 1 has been learning another language since January 2021 and did not get COVID-19 during data collection, subject 2 got COVID-19 around December 2021 while Subject 3 got COVID-19 around August 2021.

### 2.2 Data Recording

MRS data collection was embedded in a functional protocol of the BBSC project (Wang et al., 2022, 2023). This protocol includes collecting of MRS data, resting state fMRI (rs-fMRI) data, and their anatomical reference T1-weighted (T1w) MRI data, which lasts around 25 mins in total. MRS data were collected directedly after the T1w MRI and prior to the rs-fMRI scanning to avoid gradient heating effects caused by the demanding fMRI scans. MRI data were collected with a 3T MR scanner (GE Discovery MR750) with a 32-channel head coil at the Haukeland University Hospital in Bergen, Norway. The minimum reporting standards in MRS (MRSinMRS) (Lin et al., 2021) are provided in the **Supplementary Materials**, and the technical details are as follows.

Seven-minute structural T1w image data were acquired using a 3D Fast Spoiled Gradient-Recalled Echo (FSPGR) sequence with the following parameters: 188 contiguous slices acquired, with repetition time (TR) = 6.88 ms, echo time (TE) = 2.95 ms, FA (flip angle) = 12°, slice thickness = 1 mm, in-plane resolution = 1 mm × 1 mm, and field of view (FOV) = 256 mm, with an isotropic voxel size of 1 mm^3^.

After that, around four-minute ^1^H-MRS-spectra data were obtained from the ACC (voxel size 25 × 25 × 25 mm^3^) by using a single-voxel point-resolved spectroscopy (PRESS) sequence (TE/TR = 35 ms/1,500 ms, 128 repetitions). Unsuppressed water reference spectra (eight repetitions) were acquired automatically after the water-suppressed metabolite spectra. The MRS voxel is located in the ACC as illustrated in **Fig. 1**, where the center of the MNI coordinate is [2, 32, 29].

**Fig. 1.**
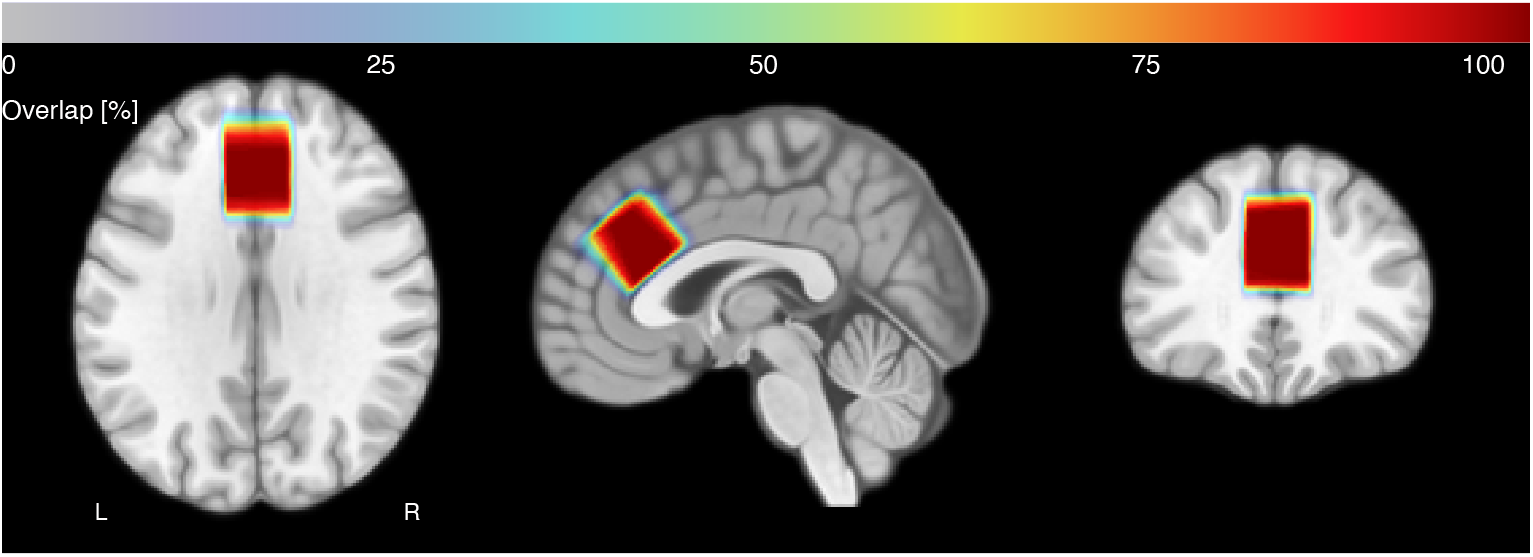
The location of the MRS voxel. The center of the MNI coordinate is [2, 32, 29]. The color spectrum represents the position overlap percentage across all sessions and all subjects. The figure was generated with Osprey 2.4.0.

### 2.3. Data Processing

The MRS data were analyzed with Osprey 2.4.0 (Oeltzschner et al., 2020) with the integrated LCModel (linear-combination model) fitting algorithm (Provencher, 1993) based on the MATLAB^®^ (R2022b) platform, which provides an automated and uniform processing pipeline including pre-processing, linear combination modeling, tissue correction, and quantification.

A concise description of the processing pipeline is as follows. The raw data were first aligned and averaged, then fitted using the LCModel which is embedded in the Osprey with the default settings across a frequency range of 0.5 to 4.0 parts per million (ppm). 22 metabolites and 9 macromolecular/lipids (**Fig. 2**) were included in the model: ascorbate(Asc), aspartate(Asp), Cr, GABA, glycerophosphocholine (GPC), glutathione (GSH), glutamine (Gln), Glu, mI, lactate(Lac), NAA, NAAG, phosphocholine (PCh), phosphocreatine (PCr), phosphorylethanolamine (PE), scyllo-inositol (sI), taurine (Tau), -CrCH2, tCho (PCh+GPC), tCr (Cr+PCr), tNAA(NAA+NAAG), and Glx (Glu+Gln).

**Fig. 2.**
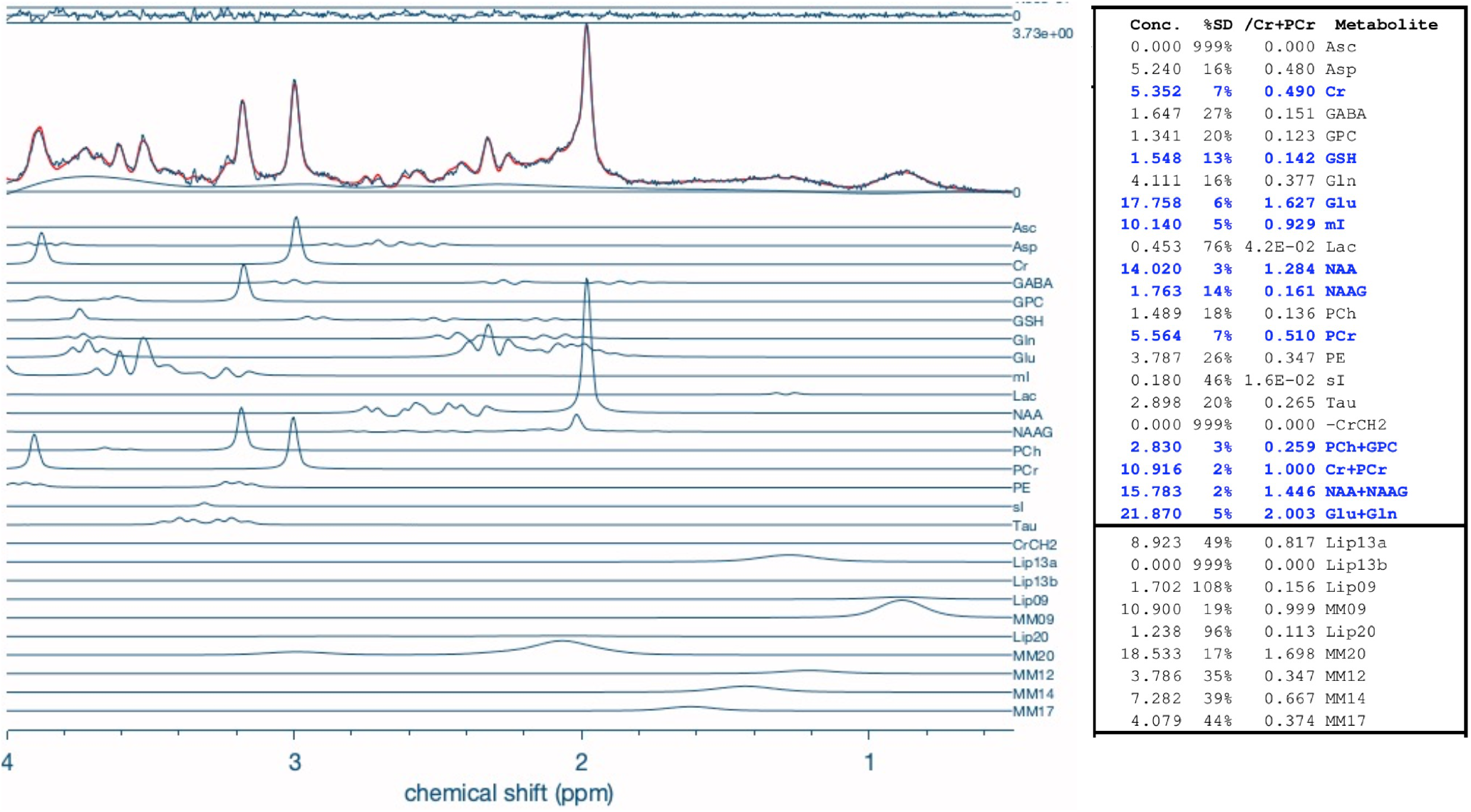
The LCModel fitting used in the Osprey. The left panel shows the metabolites used in the fitting model; the right panel shows an example result of the model fitting, where the metabolites in blue passed the quality check (criteria %SD < 15%). The lower, the better regarding the values of SD.

Before quantification, the brain was segmented into GM, WM, and cerebrospinal fluid (CSF) after coregistration to the structural image with functions from SPM12 (Friston et al., 1994) invoked by Osprey. Quantification of the metabolites was calculated using two different references: as the ratio to tCr and as water-scaled metabolite estimates with fully tissue-and-relaxation-corrected molal concentration estimates (Gasparovic et al., 2006). For more details, we recommend interested readers to the original article about Osprey (Oeltzschner et al., 2020).

Those metabolisms in blue (**Fig. 2**) have passed the quality check (%SD < 15%; the lower the better). But only those that consistently under 15% across all sessions will be reported in this study, which are Cr, PCr, tCr; Glu, Glx; NAA, tNAA; mI, tCho.

## 2.3. Statistical Analysis

To evaluate the within-subject reliability of MRS measurements, the coefficient of variance (CV) is used:

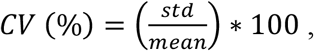

where *std* represents the standard deviation of the sample between sessions within each subject while the *mean* is the average value of the sample between sessions within each subject.

The coefficient of variation can provide the comparison of the variability of data sets with different units of measurement or scales, which has been ubiquitously used in evaluating the reliability of the MRS measurement (Baeshen et al., 2020; Brix et al., 2017; Duda et al., 2021; Kirov et al., 2012; Mikkelsen et al., 2016; Mullins et al., 2003; Near et al., 2014; O’Gorman et al., 2011; Schirmer & Auer, 2000; van Veenendaal et al., 2018). A lower CV represents a lower degree of variability relative to the mean, which indicates lower relative variability and thus greater precision or consistency in the data; while a higher CV represents a higher degree of variability relative to the mean, which indicates higher relative variability and thus lower precision or consistency.

To compare the reliability levels qualified by the ratio to tCr and water, a one-tailed paired t-test was executed since we expected that the ratio to water will generate smaller CV values.

To assess the time-of-day effect, we first separate the dataset into morning and afternoon sessions according to the recording time. In total, there are 22 morning sessions and 16 afternoon sessions in Sub1 (Sub2: 23, 16; Sub3: 15, 10). Then t-test was used to assess the significant difference between these two sessions for each subject. To control multiple comparisons, FDR correction was used. All statistical analyses were done on R 4.2.1 (R Core Team, 2022).

## 3. Results

The signal-to-noise ratio (SNR) and the linewidth of the dataset are 82 ± 11 and 5.85 ± 0.82 Hz, respectively.

### 3.1 Coefficients of variability

The scatter plots of the metabolites are illustrated in **Fig. 3**. The CVs of the metabolites in the ACC of subjects are shown in **Table 2**.

**Table 2.** The CVs of metabolites levels in the anterior cingulate cortex

|  | tCr |  |  | Water |  |  |
| --- | --- | --- | --- | --- | --- | --- |
|  | Sub1 | Sub2 | Sub3 | Sub1 | Sub2 | Sub3 |
| tNAA | 3.74 | 3.79 | 4.46 | 2.64 | 1.61 | 1.96 |
| NAA | 3.84 | 3.71 | 4.90 | 3.38 | 1.99 | 3.61 |
| tCr | N/A | N/A | N/A | 3.63 | 3.30 | 4.43 |
| PCr | 15.89 | 21.35 | 20.76 | 17.25 | 20.47 | 23.00 |
| Cr | 15.13 | 17.78 | 23.16 | 15.06 | 19.68 | 22.83 |
| Glx | 7.81 | 6.81 | 7.66 | 6.31 | 5.39 | 5.29 |
| Glu | 7.47 | 5.10 | 5.65 | 6.08 | 4.26 | 5.24 |
| mI | 7.20 | 7.72 | 7.24 | 6.49 | 6.39 | 5.37 |
| tCho | 7.27 | 7.61 | 7.12 | 6.78 | 6.41 | 7.89 |

All values are in percentage %. The smaller the better.

**Fig. 3.**
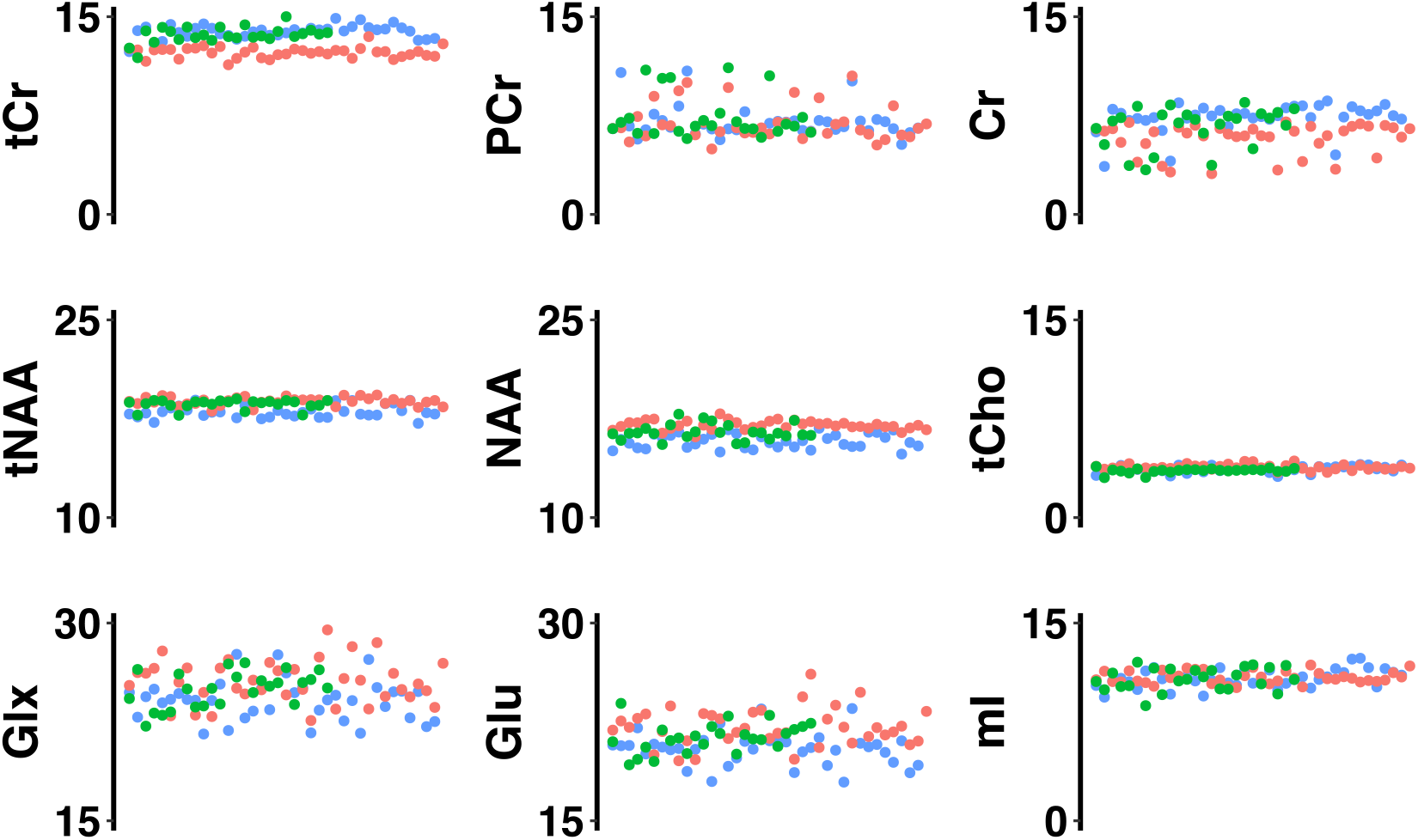
The fluctuation of the metabolites across a year. The y-axis represents the metabolite ratio to water while the x-axis represents sessions. The unit of the y-axis is an arbitrary unit (a.u.). The colors red, blue, and green represent Sub1, Sub2, and Sub3, respectively.

It is shown that the variabilities of NAA, tNAA, and tCr are low with CVs ranging from 1.61 to 4.90. In addition, the variabilities of Glu, Glx, mI, and tCho are modest with CVs ranging from 4.26 to 7.89. Furthermore, the variabilities of Cr and PCr are high with CVs between 15.06 and 23.16. Lastly, compared with the measurement of the ratio to tCr, the CVs of the water measurement are smaller (*t*_*23*_ = -2.9, *p* = 0.004, Cohen’s d = 0.59).

### 3.3 Time of Day

The levels of metabolites quantified with the ratio to water were used to examine the time-of-day effect since it showed smaller CVs. It is shown that the majority of the metabolites analyzed in this study manifested higher levels in afternoon sessions than that of morning while only PCr and NAA showed an opposite pattern (**Tables 3-5**). However, we did not find any significant difference between morning and afternoon sessions within the subject after FDR correction (**Tables 3-5**). The distributions of the metabolites of the morning and afternoon sessions are illustrated in **Fig. 4**, which were generated with the R package *raincloudplots* v0.2 (Allen, Poggiali, Whitaker, Marshall, & Kievit, 2019).

**Table 3.**
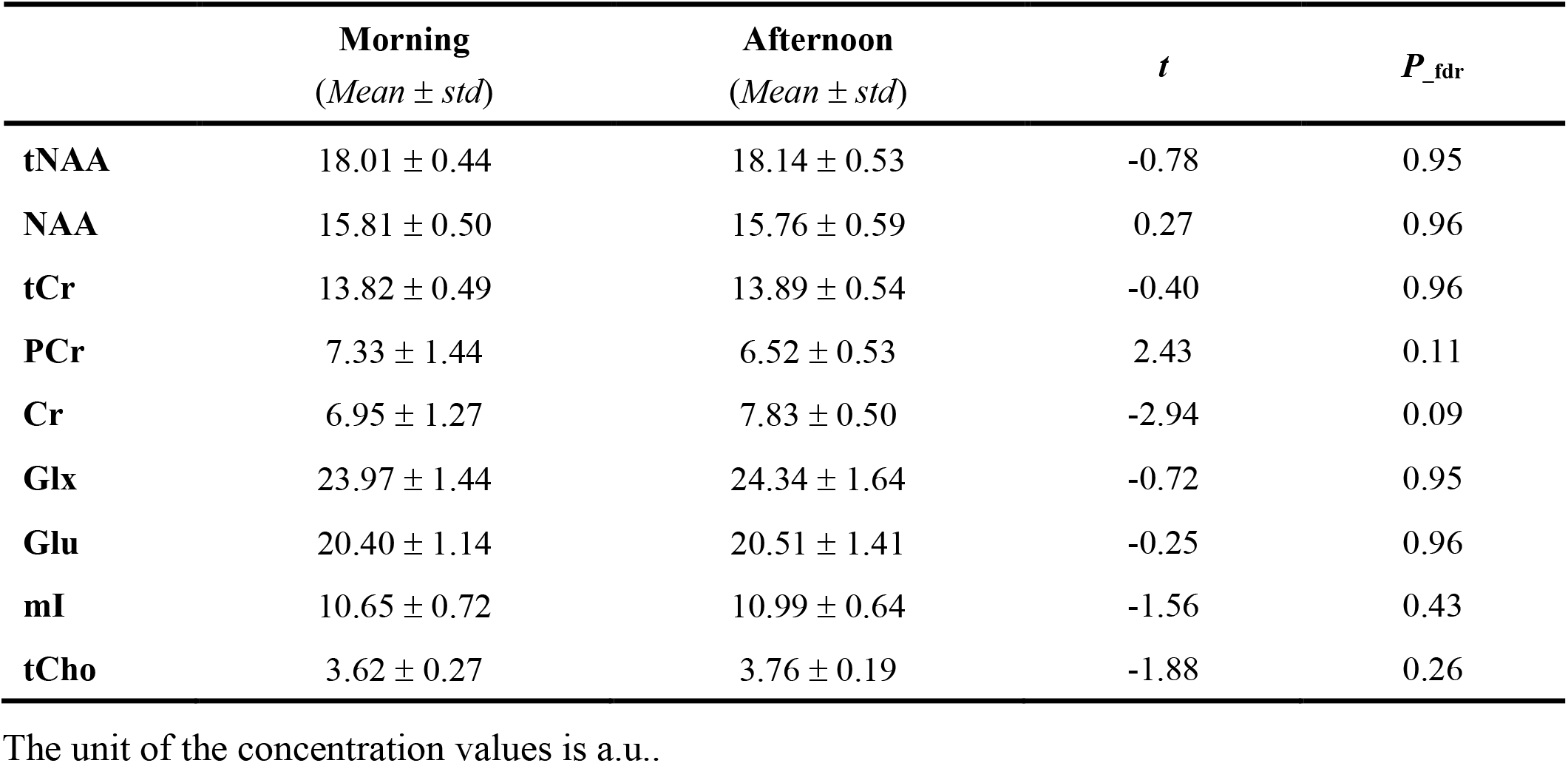
The different levels of metabolites during morning and afternoon with the ratio to water in Sub1.

**Table 4.**
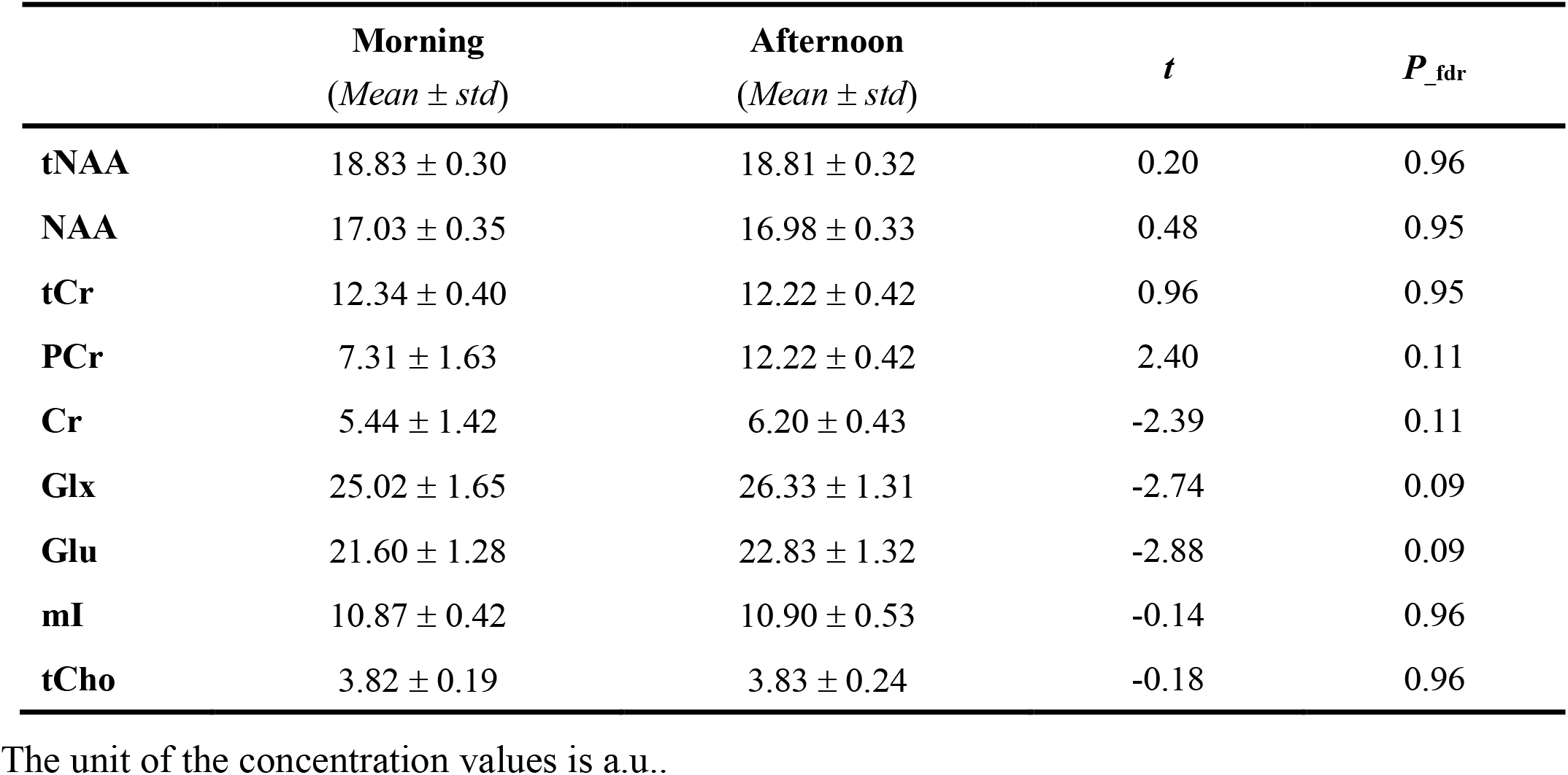
The different levels of metabolites during morning and afternoon with the ratio to water in Sub2.

**Table 5.**
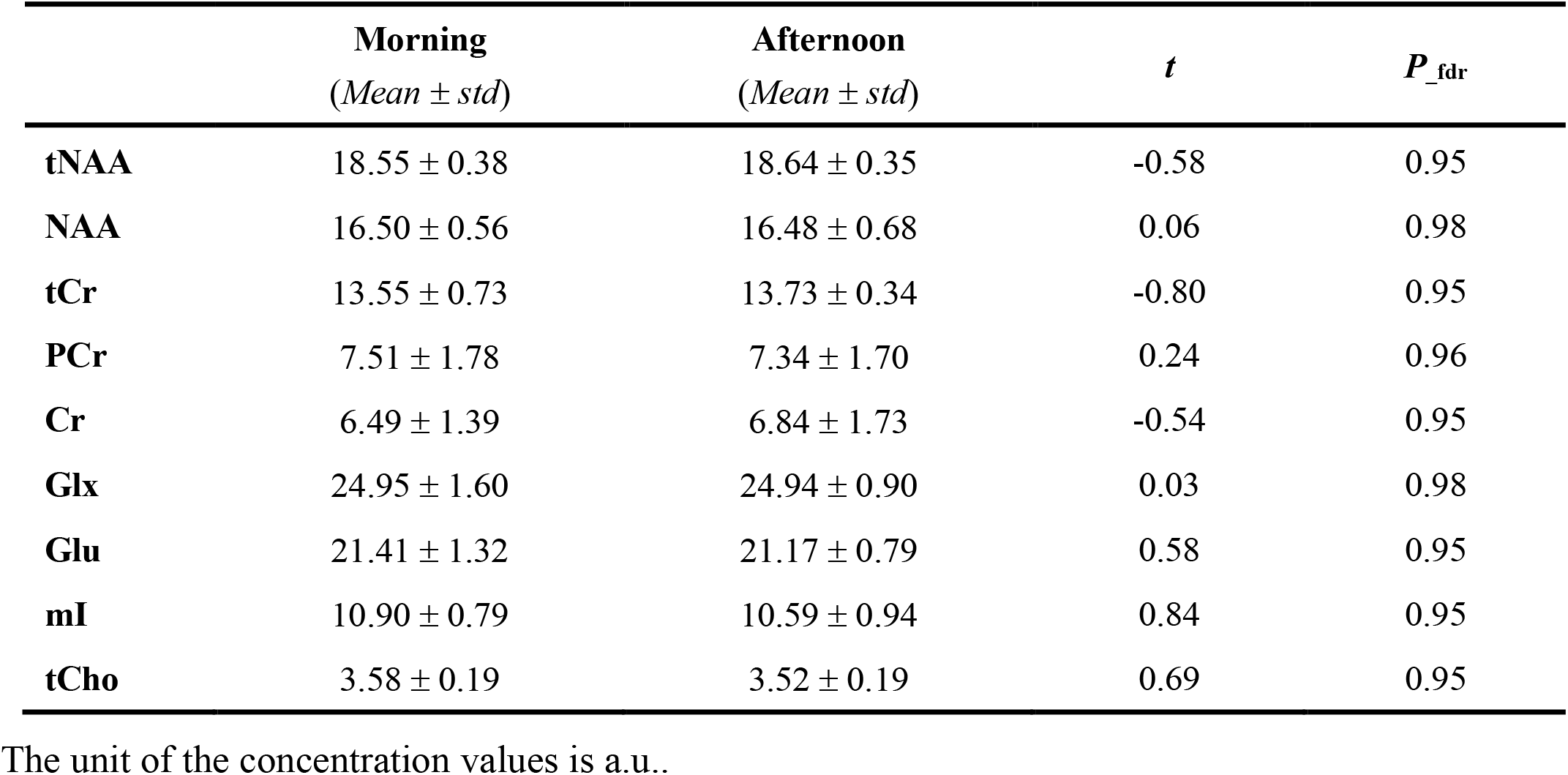
The different levels of metabolites during morning and afternoon with the ratio to water in Sub3.

**Fig. 4.**
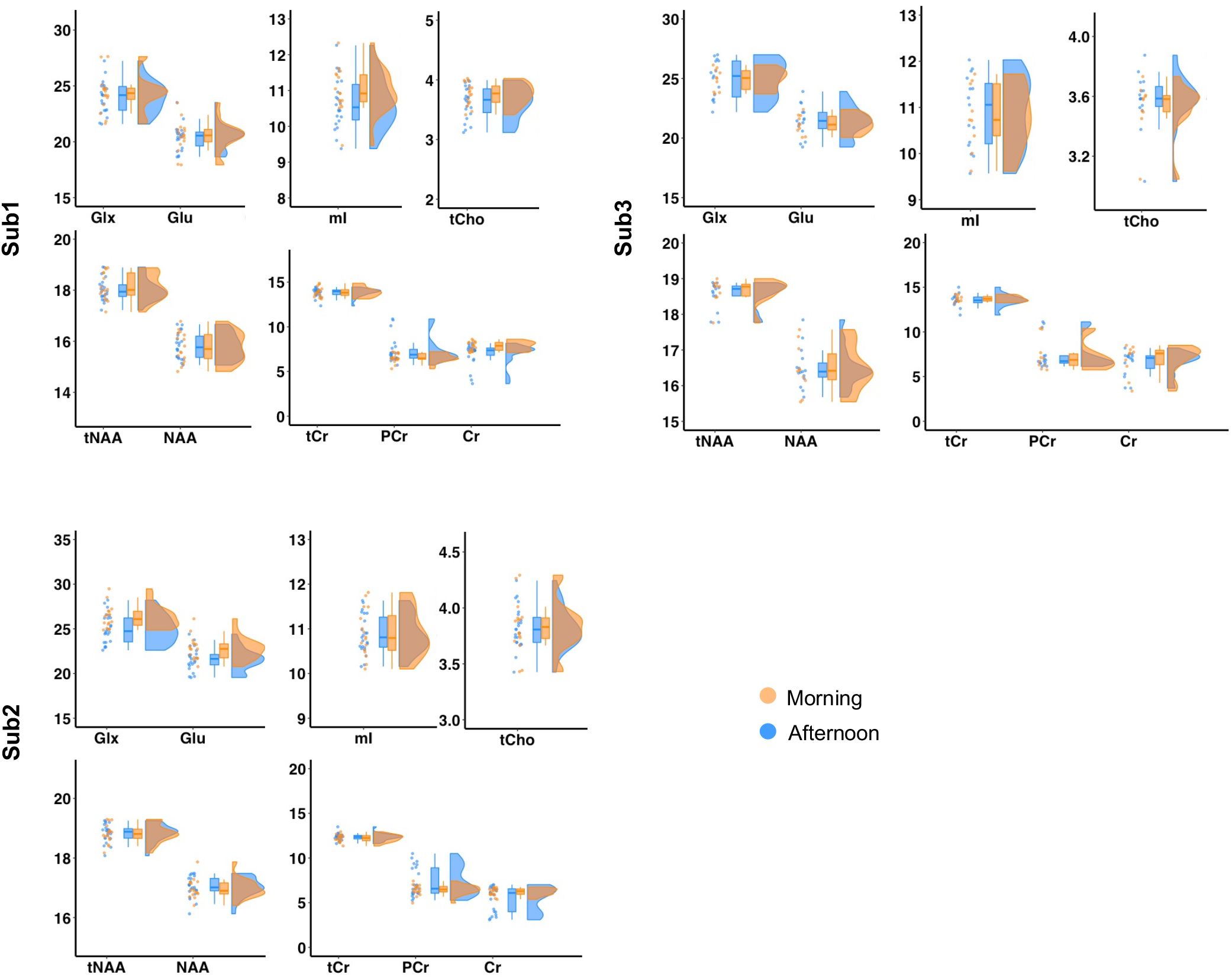
The distributions of the metabolites. The unit of the y-axis is an arbitrary unit (a.u.) and the values were calculated as the ratio to water with tissue correction.

## 4. Discussion

In this study, a longitudinal MRS dataset is constructed, where three participants were repeatedly scanned for a year with an even interval. With the aim of exploring the reliability of MRI measurements, this study specifically for the MRS measurements belongs to the BBSC Project (Korbmacher et al., 2023; Wang et al., 2022, 2023). There are three major findings. First, it is found that MRS measurements showed decent reliability with CVs ranging from 1.61% to 7.89% except for Cr and PCr, which showed lower reliability with CVs around 18%. Second, it is shown that using water as an internal concentration reference resulted in smaller CVs compared to the ratio to tCr. Third, we did not find any significant results of the time-of-day effect on MRS measurements. The potential applications of the results are discussed as follows.

### 4.1 Variability of metabolites level

As indicated by **Table** 2, we found that tNAA, NAA, Glx, Glu, tCho, and mI in the ACC showed good reliability across a year with CVs ranging from 2∼8%, which is comparable to the CVs generated from previous studies (Baeshen et al., 2020; Kirov et al., 2012; O’Gorman et al., 2011; van Veenendaal et al., 2018). To be noticed, the data collected in previous studies vary on the collection duration, MRI scanner, and brain regions. For example, the data were collected on the same day (O’Gorman et al., 2011; van Veenendaal et al., 2018), within a week (Baeshen et al., 2020), or across 3 years (Kirov et al., 2012). In conjunction with our results, it indicates that the level of metabolites is reasonably reliable across different timescales.

As expected, the level of tCr, which is the combination of Cr and PCr, showed greater reliability than Cr and PCr alone, since the constituent components are highly overlapping and difficult to disentangle from a regular PRESS sequence. Accordingly, we suggest that tCr should be emphasized instead of focusing on Cr and PCr individually.

### 4.2 The ratio to water

There are several ways to quantify the level of metabolites, of which the ratio to water and tCr are the most widely used ones. Previous studies have shown that the ratio to water performs better than that of the ratio to tCr regarding the reliability of GABA (Duda et al., 2021). Our results are in line with this finding, where the ratio to water is also better performed for other metabolites. Indeed, we found that a dominant number of previous studies, which investigate the reliability of the level of MRS measurement, used the ratio to water to quantify the metabolites level (Baeshen et al., 2020; Bogner et al., 2010; Brix et al., 2017; Duda et al., 2021; Mikkelsen et al., 2016; O’Gorman et al., 2011; van Veenendaal et al., 2018) instead of the ratio to tCr as the internal concentration reference (Near et al., 2014; Schirmer & Auer, 2000). More importantly, a recent consensus recommendation suggests using the ratio of water (Near et al., 2021). Accordingly, we advocate the ratio to water as the internal concentration reference in MRS studies.

### 4.3 The time-of-day effect

Despite there being evidence showing that different data collection time points could affect the functional brain organizations (Orban et al., 2020; Vaisvilaite et al., 2022), we did not find this effect on the levels of metabolites. Correspondingly, it corroborates the indication that the MRS measurement is reasonably stable.

### 4.4 Limitations

Before making any conclusions, some limitations should be articulated. First, phantom data can be used to assess the absolute estimate of metabolite concentrations and indicate the reliable performance of the MRI scanner (Brix et al., 2017; van Veenendaal et al., 2018).

Since we found that the MRS measurement is reasonably reliable, we believe it will be much more stable when ruling out the variance contributions from the measurement itself. Second, only the default LCM algorithm was utilized which could affect the conclusion, since different LCM algorithms could provide different quantifications (Zollner et al., 2021). Third, the sample size is relatively small compared to other conventional studies. However, the present new data collection method, where the same participants are scanned repeatedly for a period of time, doesn’t necessarily need a very large sample. But it can provide invaluable insights that conventional studies cannot offer (Korbmacher et al., 2023; Wang et al., 2022, 2023). Lastly, the absence of female participants may affect the conclusions of this study, since there is evidence suggesting that the menstrual cycle could influence the GABA level (Harada, Kubo, Nose, Nishitani, & Matsuda, 2011) and other metabolites level (Hjelmervik et al., 2018).

## 5 Conclusion

In summary, it is shown that the MRS measurement is quite reliable in detecting the concentration of the metabolites even across a year. In addition, the ratio to water is suggested to use as the internal concentration reference.

## Supporting information

The minimum reporting standards in MRS (Lin et al., 2021) are provided in the Supplementary Materials, and the technical details are as follows.

## Data and code

Data and code used in this manuscript can be found here on Github https://github.com/MengYunWang/BBSC/tree/main/MRS.

## Acknowledgments

We appreciate the technical support and data collection support from our radiologists (Christel Jansen, Eva Øksnes, Roger Barndon, Trond Øvreaas, Tor Fjørtoft, and Turid Randa) at the Haukeland University Hospital.

