## Supplementary material for "The intra-individual reliability of ^1^H-MRS measurement in the anterior cingulate cortex across one year": The minimum reporting standards in MRS (Lin et al., 2021) are provided in the Supplementary Materials, and the technical details are as follows.

**Table 1**. MRSinMRS checklist.

| Site (Name or Number) |  |
| --- | --- |
| 1. Hardware |  |
| a. Field strength [T] | 3T |
| b. Manufacturer | GE |
| c. Model (software version if available) | Discovery MR750 |
| d. RF coils: nuclei (transmit/ receive), number of channels, type, body part | 32-channel head coil |
| e. Additional hardware |  |
| 2. Acquisition |  |
| a. Pulse sequence | single-voxel point-resolved spectroscopy (PRESS) sequence |
| b. Volume of Interest (VOI) locations | Anterior Cingulate Cortex |
| c. Nominal VOI size [cm^3^, mm^3^] | 25 x 25 x 25 mm3 |
| d. Repetition Time (TR), Echo Time (TE) [ms, s] | TR 1500 ms, TE 35 ms |
| e. Total number of Excitations or acquisitions per spectrum  In time series for kinetic studies   i. Number of Averaged spectra (NA) per time-point ii. Averaging method (e.g. block-wise or moving average) iii. Total number of spectra (acquired / in time-series) | 16 total averages with 16 averages per subspectrum |
| f. Additional sequence parameters  (spectral width in Hz, number of spectral points, frequency offsets)  If STEAM:, Mixing Time (TM)  If MRSI: 2D or 3D, FOV in all directions, matrix size, acceleration factors, sampling method | F1: 5000 Hz, 4096 points |
| g. Water Suppression Method | CHemical Shift Selective saturation (CHESS) |
| h. Shimming Method, reference peak, and thresholds for “acceptance of shim” chosen | Vendor default prescan (double-echo GRE) |
| i. Triggering or motion correction method  (respiratory, peripheral, cardiac triggering, incl. device used and delays) | n/a |
| 3. Data analysis methods and outputs |  |
| a. Analysis software | Osprey 2.4.0 |
| b. Processing steps deviating from quoted reference or product | None |
| c. Output measure  (e.g. absolute concentration, institutional units, ratio)Processing steps deviating from quoted reference or product | tCr, Tissue Correted Water Scaled |
| d. Quantification references and assumptions, fitting model assumptions | LCModel basline knot spacing 0.40 ppm |
| 4. Data Quality |  |
| a. Reported variables  (SNR, Linewidth (with reference peaks)) | SNR (Cr): SNR: 82 ± 11 Hz  linewidth (Cr): linewidth 5.85 ± 0.82 Hz |
| b. Data exclusion criteria | SD < 15% |
| c. Quality measures of postprocessing Model fitting (e.g. CRLB, goodness of fit, SD of residual) | SD of residual |
| d. Sample Spectrum | Figure 2 |

Lin, A. D., Andronesi, O., Bogner, W., Choi, I. Y., Coello, E., Cudalbu, C., . . . Sta, E. W. G. R. (2021). Minimum Reporting Standards for in vivo Magnetic Resonance Spectroscopy (MRSinMRS): Experts' consensus recommendations. NMR in Biomedicine, 34(5). doi:10.1002/nbm.4484
